# Differential variation and expression analysis

**DOI:** 10.1101/276337

**Authors:** Haim Bar, Elizabeth D. Schifano

## Abstract

We propose an empirical Bayes approach using a three-component mixture model, the *L*_2_*N* model, that may be applied to detect both differential expression (mean) and variation. It consists of two log-normal components (*L*_2_) for the differentially expressed (dispersed) features (one component for under-expressed [dispersed] and the other for over-expressed [dispersed] features), and a single normal component (*N*) for the null features (i.e., non-differentially expressed [dispersed] features). Simulation results show that *L*_2_*N* can capture asymmetries in the numbers of over-and under-expressed (dispersed) features (e.g., genes) when they exist, can provide a better fit to data in which the distributions of the null and non-null features are not well-separated, but can also perform well under symmetry and separation. Thus the *L*_2_*N* model is particularly appealing when no a priori biological knowledge about the mixture density is available. The *L*_2_*N* model is implemented in an R package called DVX, for **D**ifferential **V**ariation and e**X**pression analysis. The package also includes an implementation of differential expression analysis via the limma package, and a differential variation and expression analysis using a three-way normal mixture model. DVX is a user-friendly, graphical interface implemented via the ‘Shiny’ package [6], so that a user is not required to have R programming knowledge. It offers a set of diagnostics plots, data transformation tools, and report generation in Microsoft Excel- and Word-compatible formats. The package is available on the web, at https://haim-bar.uconn.edu/software/DVX/.

## 1 Introduction

High-dimensional data arise frequently in health sciences and biomedical studies, and has emerged in recent years as a consequence of the rapid advance of “-omic” research. For example, array-based technologies allow scientists to simultaneously collect measurements on hundreds of thousands of genetic markers from an experimental sample. Due to high cost, it is common that these markers are measured for a relatively small number of independent samples in a given study, and as a consequence, one faces the ‘large *G*, small *n*’ problem, where *G* is the total number of markers or features and *n* is the number of samples. The resulting datasets consist of observed values quantifying relative abundance levels of platform-dependent biological material at multiple sites across the genome. Particularly for gene expression and DNA methylation array experiments, the goal is often to identify genetic markers that are differentially expressed (methylated) across clinically relevant subgroups. Similar settings and goals are also found in next-generation sequencing platforms (e.g., RNA-seq), as well as fields beyond genomics (e.g., metabolomics, brain imaging), however, the terminology of gene expression will be used below for simplicity of exposition. As such, the objective is to conduct simultaneous tests across the *G* genetic markers for differential mean expression due to a particular predictor variable of interest (e.g., clinical subgroup, age group, etc).

The existing methods for detecting differential mean expression between two populations are numerous, as classical parametric (t-/F-) statistics evaluated at each marker do not provide a reliable methodology for determining differential mean expression across the genome. The large number of markers with relatively few samples not only induces a severe multiple testing problem, but also yields marker-wise variance estimates that are often inaccurate [e.g. 20, 8]. More powerful tests have since been proposed that combine information across genetic markers for stabilization. Indeed, the most widely-used methods currently for detecting differential gene expression ‘borrow strength’ across genetic markers by treating marker-specific effects as random variates [e.g., 18, 7, 10, 1, 3].

In this work, we take a broader interpretation of ‘differential expression’ and allow for identification of genes which are differential across populations not only in terms of their mean expression levels, but also possibly their variances. Differential variation is important, for example, in the analysis of heritability of complex diseases [16], epigenetic analysis [9], and gene network regulation [17]. Recently, Bar et al. [2] proposed an extension of the ‘lemma’ model in [1, 3] by introducing a bivariate modeling strategy which accounts for both differential mean expression and differential dispersion across two populations, and yields a substantial gain in power to detect differential mean expression when differential dispersion is present. As with the limma [18] and lemma [1] models, Bar et al. [2] uses a mixture model, but in contrast to limma and lemma, it is based on a mixture of three normal distributions with one component designed to capture the non-differentially expressed (dispersed) genes, and the remaining two components designed to capture the underexpressed (underdispersed) and overexpressed (overdispersed) genes relative to a reference group. The two-component mixture models of limma [18] and lemma [1] implicitly assume that the differentially expressed genes are symmetrically up- and down-regulated, which may or may not be reasonable depending on the data at hand (see, e.g., [11]). In the extreme case where only up- or down-regulated genes are expected or relevant, Ivanek et al. [11] proposed first using limma [18] to estimate certain hyperparameters and rank the genes, and then fitting an extreme value distribution to the tail of interest. While the three-component Normal mixture of Bar et al. [2] can adequately capture asymmetries in the numbers of over- and under-expressed (dispersed) genes when they exist, the performance of the three-component Normal mixture model can be suboptimal when there is a large degree of overlap between the three mixture components. In this situation, for example, a large portion of the density near zero may be allocated counterintuitively to the nonnull components, the mixture component corresponding to the over-expressed genes may attribute an overly large probability to negative values of test statistic (e.g., difference in means), and vice versa.

To overcome these challenges, we propose an empirical Bayes approach using a different three-component mixture model, the *L*_2_*N* model, that consists of two log-normal components (*L*_2_) for the differentially expressed (dispersed) genes (one component for under-expressed (dispersed) and the other for over-expressed (dispersed) genes), and a single normal component (*N*) for the null genes (i.e., non-differentially expressed (dispersed) genes). Our simulation results show that this approach can still capture asymmetries in the numbers of over- and underexpressed (dispersed) genes when they exist, but can also provide a better fit to data exhibiting a large degree of overlap in the three-component Normal mixture while still providing a good fit when the numbers of over- and underexpressed (dispersed) genes are similar. Thus the *L*_2_*N* model is particularly appealing when no a priori biological knowledge about the mixture density is available. This type of mixture, in which the non-null components have zero mass or density at the center of the null distribution and a negligible mass in a sufficiently small neighborhood around it, is similar to what was defined in [12] as ‘non-local alternative priors’ in the Bayesian hypothesis testing framework. In that paper, Johnson and Rossell use point-mass null and a different form of alternative which, unlike the *L*_2_*N* model, is symmetric and assigns equal probabilities to over- and underexpressed genes. They further show (under additional constraints) the gain that their model yields in terms of rate of convergence of the Bayes factor.

The *L*_2_*N* model is implemented in an R package called DVX [4], for **D**ifferential **V**ariation and e**X**pression analysis. The package also includes an implementation of a three-way normal mixture model, which we refer to as *N*_3_, similar to the one proposed in [2]. The software also allows for differential expression analysis via the limma package. DVX is a user-friendly, graphical interface implemented via the ‘Shiny’ package [6], so that a user is not required to have R programming knowledge. It offers a set of diagnostics plots, data transformation tools, and report generation in Microsoft Excel- and Word-compatible formats. All three models implemented in DVX (namely, *L*_2_*N, N*_3_, and limma) allow for adjustment for covariates. Furthermore, DVX allows users to define more general contrasts than the simple two-group comparison. An extensive documentation of the software is provided in the Supplementary Materials.

The paper is organized as follows. We introduce the *L*_2_*N* model in Section 2 and further describe the relationship between the three models implemented in DVX. In Section 3 we present results from the three model implementations under various simulation experiments as well as a case-study in which we identify change in gene expression levels as a result of aging (across four age groups) in normal brains [15]. We conclude with a brief discussion in Section 4.

## 2 Methods

We are interested in identifying which genes have different distributional properties of expression levels between two populations. With normalized expression data this is usually interpreted as testing for differences between the means of two distributions. However, we use a more general interpretation and ask which genes are differentially expressed (different means) and/or differentially dispersed (different variances) between the two populations.

In the following, the normalized expression levels for gene *g* in group *i* = 1, 2 have mean *μ*_*ig*_ and variance 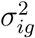. The sample versions are *m*_*ig*_ and 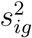, respectively. A gene is differentially expressed (DE) if *β*_*g*_ = *μ*_1*g*_ *- μ*_2*g*_ =*≠* 0. With normalized data, testing the null hypotheses *H*_0*g*_: *β*_*g*_ = *θ*_0_ relies on a statistic of the form

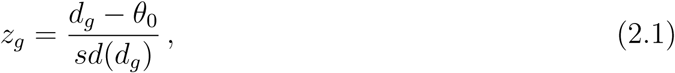

where *d*_*g*_ = *m*_1*g*_ *- m*_2*g*_. For non-DE genes, if *var*(*d*_*g*_) are assumed to be known, *z*_*g*_ has a standard normal distribution for non-DE genes. Otherwise, *z*_*g*_ follows a *t* distribution. A common approach to detecting DE genes is to use a mixture model in which non-DE genes are assumed to follow a ‘null distribution’ and belong to one component, and DE genes follow another distribution and belong to a different mixture component. For example, in the limma model [18], the variances are assumed equal across groups, i.e., 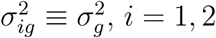, and 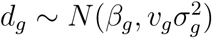 such that for non-DE genes *β*_*g*_ = 0 and for DE genes 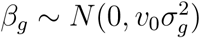. This results in a two-component mixture model for 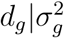, in which both components have mean zero, but expression levels of genes in the DE component have greater variability. The model in [2] is a three component mixture in which for non-DE genes 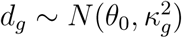, where 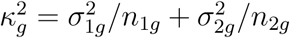, and for DE genes 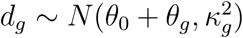 where *θ*_*g*_ *∼ N* (*±θ*_*D*_, *κ*^2^). This model consists of three normal components, with means *θ*_0_, *θ*_0_ *- θ*_*D*_, and *θ*_0_ + *θ*_*D*_, where *θ*_*D*_ *>* 0. Here, we use a more general model in which the means of the non-null components are *θ*_0_ + *θ*_*D*1_ and *θ*_0_ *- θ*_*D*2_ and where *θ*_*D*1_ may be different than *θ*_*D*2_. We denote this model by *N*_3_. Table 1 summarizes the differences between three approaches for detecting DE: the standard *t*-test (one gene at a time), limma, and *N*_3_. A key difference between the three models is in how *var*(*d*_*g*_) is estimated. In the standard *t*-test approach the variances are estimated separately for each gene and each group, while in the mixture models (limma and *N*_3_) the variances are estimated by borrowing information across all genes. This is achieved by assuming a prior distribution for the error variances of all genes. Typical gene expression experiments involve small sample sizes, and thus lack power to detect differentially expressed genes. Power is further reduced when one accounts for multiple testing and adjusts the significance level for the large number of hypotheses (one per gene). When sample sizes are small (or even moderate) and the number of genes is large, the mixture models yield a substantial increase in power for detection of DE genes, due to the so-called ‘shrinkage estimation’ [e.g., 18, 10].

**Table 1:** Models for differential expression. *β*_*g*_ = *μ*_1*g*_ *- μ*_2*g*_, *d*_*g*_ = *m*_1*g*_ *- m*_2*g*_. In all three models inference is based on *z*_*g*_ from (2.1).

| Model | $\beta_g\vert H_0$ | $\beta_g\vert H_1$ | $var(d_g)$ |
| --- | --- | --- | --- |
| $t$-test | $\theta_0 = 0$ | $\neq 0$ | $\frac{\sigma_{1g}^2}{n_{1g}} + \frac{\sigma_{2g}^2}{n_{2g}}$ |
| limma | $\theta_0 = 0$ | $\sim N(0, v_0\sigma_g^2)$ | $\sigma_g^2 \left( \frac{1}{n_{1g}} + \frac{1}{n_{2g}} \right)$ |
| $N_3$ | $\theta_0$ | $\sim N(\theta_0 + \theta_{D1}, \kappa^2)$ if $\beta_g > \theta_0$<br>$\sim N(\theta_0 - \theta_{D2}, \kappa^2)$ if $\beta_g < \theta_0$ | $\frac{\sigma_{1g}^2}{n_{1g}} + \frac{\sigma_{2g}^2}{n_{2g}}$ |

**The** *L*_2_*N* **Model**: In the model presented in this paper we also assume that {*z*_*g*_} come from a mixture distribution, and that non-DE genes follow a normal distribution, 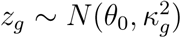, where 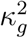 are also assumed to follow a prior distribution. For the DE genes, we assume that

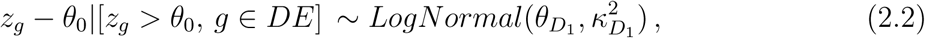

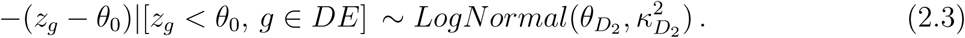

The parameter *θ*_0_ represents an overall difference between non-DE genes in the two groups. While it may be close to 0 in many applications, this is not always the case, and models which assume that *θ*_0_ = 0 when it is not so, yield many false discoveries as we discuss in Section 3.3.

Denote the non-DE probability density function (p.d.f.) of *z*_*g*_ by *f*_0_ and the p.d.f.s of the two DE components by *f*_1_ and *f*_2_. Denote the three components of the mixture by *C*_0_, *C*_1_, or *C*_2_, and let the corresponding proportions of genes belonging to each component *j* = 0, 1, 2, be *p*_*j*_ such that 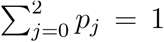. Classifying genes into one of the three components is then achieved by computing their posterior probabilities

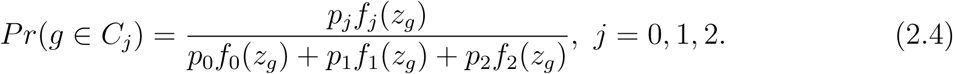

This mixture model, which we call *L*_2_*N*, allows for different proportions of overexpressed and underexpressed genes. Under this model, the probability that a gene with a positive (negative) *z*_*g*_ statistic is misclassified as underexpressed (overexpressed) is zero. In contrast, the limma model assumes that the distribution of DE genes is symmetric, and while the *N*_3_ model allows for different proportions of over and underexpressed genes, the unbounded support of the DE components implies that an overexpressed gene has a non-zero probability of being classified as underexpressed, and vice versa. Figure 1 demonstrates the three types of mixtures mentioned here, with the limma model on the left, the *N*_3_ model in the middle, and the *L*_2_*N* model on the right. The dotted blue lines represent the distributions of the non-DE genes, which are distributed normally, and in all three models we set *Pr*(*g* ∈ *C*_0_) = 0.8. The dashed red lines represent the distributions of DE genes, and the thick gray lines represent the mixture distributions. In the limma model, the assumptions imply symmetry, and thus, that the proportions of overexpressed and underexpressed genes are the same. This may be a biologically reasonable assumption in some situations, but not in others. In both the limma and *N*_3_ models, the probability of a type II error is greater than in the *L*_2_*N* model - for a DE gene, i.e. *g* ∈ ∉ *C*_0_, and for some small *E >* 0, *P* (*|z*_*g*_*| < E | g* ∈∉ *C*_0_) is smaller for *L*_2_*N* than for the limma and *N*_3_ models.

**Figure 1:**
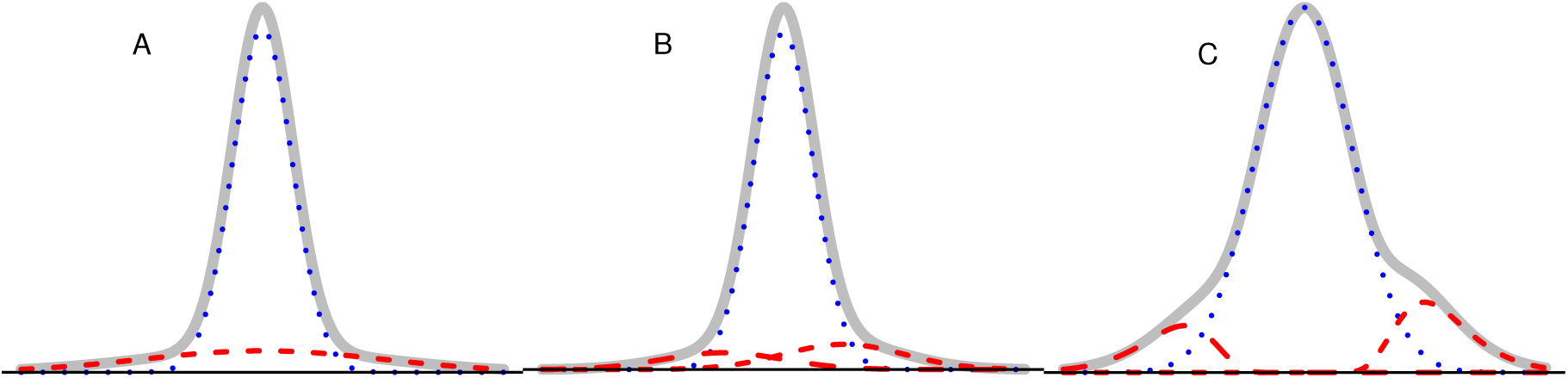
Three mixture models for differential expression. A. The limma model [18] B. the *N*_3_ model [2] C. the *L*_2_*N* model. In all cases, *Pr*(*g* ∈ *C*_0_) = 0.8.

Both the *N*_3_ and *L*_2_*N* models use a hierarchical mixed model in which 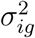, *i* = 1, 2, are assumed to have a common prior distribution for all *g*, within group *i*. This allows ‘borrowing information’ across genes and shrinks larger variances toward the overall mean of variances. To obtain ‘shrinkage estimates’ for the variances, 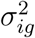, we follow [2] and define

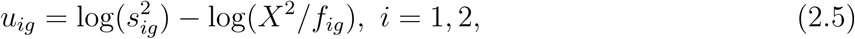

where *X*^2^ is a Chi-squared random variate with *f*_*ig*_ = *n*_*ig*_ - 1 degrees of freedom. Given 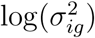,

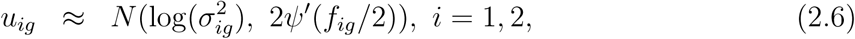

where *Ψ*^*′*^ is the trigamma function. The normal approximation in (2.6) allows us to test for differential dispersion, which may be of interest in its own right. To do that, we place a three-component *L*_2_*N* prior on 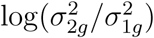. Then, the differential dispersion statistics

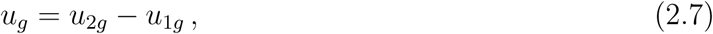

follow an *L*_2_*N* mixture distribution having the same form as (2.2) and (2.3): using the superscript *v* to indicate that the model parameters for the dispersion statistics are different from those for the expression model, for differentially dispersed (DD) genes we assume

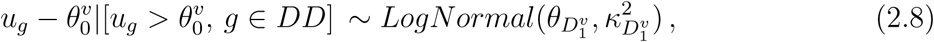

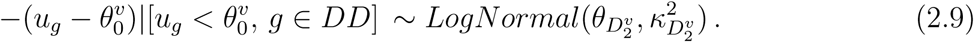

Similarly to the expression model, we denote the mixture components for *u*_*g*_, by 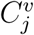 *j* = 0, 1, 2 (which may be different from the DE mixture components, *C*_*j*_), and the posterior probabilities 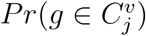, have the same form as (2.4).

We compute the posterior mean of *σ*^2^, *i* = 1, 2, denote it by 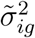 and fit the *L*_2_*N* model to {*z*_*g*_}, with 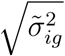 replacing the *sd*(*d*_*g*_) in (2.1). To fit the *L*_2_*N* model to {*z*_*g*_} (or {*u*_*g*_}), we use the EM algorithm, where the ‘missing data’ are indicator variables such that *b*_*gj*_ = 1 for *g* ∈ *C*_*j*_ (or 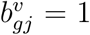 for 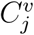), *j* = 0, 1, 2, and *b*_*gj*_ = 0 if *g* ∈∉ *C*_*j*_ (or 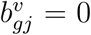 if 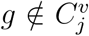). We estimate 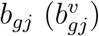 by taking its expectation (i.e., (2.4) for expression; similar for dispersion), given the current estimates of the mixture component parameters. Maximum likelihood estimates for *θ*_0_, *θ*_*D*_*j*__, and 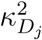 (or 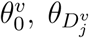, and 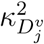) are obtained in each iteration, while holding the values *b*_*gj*_ fixed at their current estimates. Additional details regarding the estimation procedure are provided in the Supplementary Materials. If *Pr*(*g* ∈ *C*_1_ *∪ C*_2_) is sufficiently large, we conclude that gene *g* is DE. Similarly, if the posterior probability 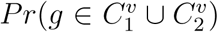 is sufficiently large, we conclude that gene *g* is DD. The normality of the null component under the *L*_2_*N* model also allows us to use the frequentist approach so that gene *g* is DE (DD) if *h*[2(1-Φ(*| z*_*g*_ |))] *< q* (or, *h*[2(1 − Φ(*|u*_*g*_|))], when testing for differential dispersion), where *h* is a function which adjusts the p-values to account for multiple testing.

## 3 Results

Experiments which aim to identify differentially expressed genes are usually performed under the assumption that the proportion of DE genes is relatively small. When the total number of genes is large and the sample sizes are small or moderate, as is often the case, the challenge from the statistical point of view is to maximize the power (i.e., find the largest number of true DE genes), while limiting the false discovery rate (FDR). The goal of the simulations presented here was to evaluate and compare the performance of the three models implemented in DVX (limma, *N*_3_, and *L*_2_*N*) in terms of power and FDR. To clarify the terminology used in this section, when we refer to the performance of limma, *N*_3_, or *L*_2_*N* we mean the software implementation of the corresponding mixture model.

In each of almost 50 different configurations we simulated 5,000 genes and *n* = 5 subjects in each group. Each configuration was simulated 30 times. We varied the number of DE genes, the mean of the difference between expression for DE genes between the two groups, which we denote by *θ*, and the variance of the random error. The error variances were simulated from an inverse gamma distribution: 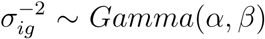. The shape and scale parameters, *α* and *β*, were set so that *E*[*V ar*(*z*_*g*_)] = 1, so that we can control the mean signal to noise ratio (SNR) in the simulation only in terms of the signal, *θ*. With this setting, *β* = *n*(*α-*1)/2, and the variance of {*V ar*(*z*_*g*_)} is determined by *α*. We let *α* ∈ {4.5, 6, 7.5, 9}, corresponding to *V ar*[*V ar*(*z*_*g*_)] ∈ {1, 0.63, 0.45, 0.36}. Here, we show results with *α* = 9. We note that as *α* increases, the variances across genes become more homogenous, and the power to detect DE genes increases for all three methods (for a fixed mean difference between the two groups, *θ*; results not shown).

With each of the three methods, the null distribution is obtained, and a gene is declared as DE if its Benjamini-Hochberg [5] adjusted p-value is less than 0.05. Note that all three methods allow for Bayesian inference, as well, in terms of posterior probabilities or Bayes’ factors.

### 3.1 Simulation Study – Differential Expression, no Differential Dispersion

First, we consider scenarios in which the data are generated according to the *N*_3_ model, when there are no differentially dispersed genes. We compare the three methods under consideration for different signal strengths, and different number of DE genes. Table 2 shows the median power of the three methods for *θ* ∈ {2.5, 3}. The median for each configuration was taken over 30 iterations. Note that the observed FDR for all three methods in the configurations described here, was indeed, on average, controlled at the desired level. In the table, *D*_1_ and *D*_2_ denote the number of genes that are over-expressed in groups 1 and 2, respectively. When *θ* and *D* = *D*_1_ + *D*_2_ are large, the three methods have similar power (defined as the fraction of the *D* differentially expressed genes correctly classified.) However, as the signal strength or the total number of DE genes decreases, *N*_3_ and *L*_2_*N* tend to have higher power than limma. Of course, the power of all methods increases with *θ*. What is perhaps less obvious is that the power decreases with *D*. The difference in power between the methods in these situations is particularly important because in many cases, it is believed that the proportion of DE genes is small. For example, when *G* = 20, 000 and *D* = 400 (proportional to the third row in Table 2), and *θ* = 3, the 4% difference in power between *L*_2_*N* and limma translates into 16 additional true DE discoveries. Arguably, from a practical standpoint, finding these additional 16 genes could have important consequences.

**Table 2:** Simulation results – median true positive DE genes. *G* =5,000 genes, *n*_1_ = *n*_2_ = 5.

| $D_1$ | $D_2$ | $\theta = 2.5$ | | | $\theta = 3$ | | |
| --- | --- | --- | --- | --- | --- | --- | --- |
| | | limma | $N_3$ | $L_2N$ | limma | $N_3$ | $L_2N$ |
| 500 | 500 | 0.31 | 0.3 | 0.31 | 0.59 | 0.57 | 0.57 |
| 225 | 25 | 0.1 | 0.13 | 0.13 | 0.35 | 0.36 | 0.35 |
| 50 | 50 | 0.05 | 0.08 | 0.08 | 0.2 | 0.22 | 0.24 |
| 35 | 15 | 0.03 | 0.06 | 0.06 | 0.14 | 0.19 | 0.19 |

We also simulated data according to the limma model, and in this scenario the three method have identical power. For example, with a total of *D* = 250 DE genes, the powers obtained by all three methods for *v*_0_ = 6, 9, 12 are 0.18, 0.28, and 0.35, respectively. When we hold *v*_0_ fixed and decrease *D* we again observe that the power decreases. For example, with *v*_0_ = 9, the powers of all three methods for *D* = 1000, 500, 200, 100 are 0.34, 0.3, 0.27, 0.24, respectively.

### 3.2 Simulation Study – Differential Expression and Differential Dispersion

We simulated datasets in which some genes were differentially dispersed across the two groups, as well as differentially expressed. The goal was to test not only whether the *L*_2_*N* method can detect those differentially dispersed genes, but also to check if and how the presence of DD genes affects the power to detect DE genes. The results presented here were obtained with *α* = 9 (relatively low variability of the error variance), and *θ* ∈ {2.5, 3}. We simulated 50 over-dispersed genes in group 1 and 50 over-dispersed genes in group 2, by dividing the standard deviation for the DD genes in one of the groups by 6.

Table 3 shows the results for *D* = 500 and *D* = 250, for *θ* = 2.5, 3. It is clear that the approaches of *N*_3_ and *L*_2_*N*, where the variance estimates are obtained from a threeway mixture model for differential dispersion, greatly increase the power to detect DE genes when DD is present as compared with limma. We also see that *L*_2_*N* is slightly more powerful than *N*_3_. When the variance of the null distribution increases (i.e., when *α* decreases) the difference in power between *L*_2_*N* and *N*_3_ increases. For example, with *α* = 6, *θ* = 2.5, and 250 DE genes, *L*_2_*N* has a power of 0.46, vs. 0.44 for *N*_3_ (and limma has a power of 0.15).

**Table 3:** Simulation results – median true positive DE genes when 100 genes are also differentially dispersed. *G* =5,000 genes, *n*_1_ = *n*_2_ = 5.

| $D_1$ | $D_2$ | $\theta = 2.5$ | | | $\theta = 3$ | | |
| --- | --- | --- | --- | --- | --- | --- | --- |
| | | limma | $N_3$ | $L_2N$ | limma | $N_3$ | $L_2N$ |
| 250 | 250 | 0.16 | 0.35 | 0.36 | 0.38 | 0.55 | 0.55 |
| 200 | 50 | 0.18 | 0.45 | 0.46 | 0.35 | 0.60 | 0.61 |

As was the case in the previous subsection, the observed FDR for all three methods was controlled at the desired level.

### 3.3 Simulation Study – Mean Shift

The three-way mixture models, *N*_3_ and *L*_2_*N*, include a term for an overall difference between the mean expression in the two groups, whereas the limma model assumes there is none. This difference was denoted earlier by *θ*_0_. To understand why it is important to account for this difference, we also simulated data sets in which *θ*_0_ = 0.5 and *θ*_0_ = 1. In these simulations no genes were differentially dispersed.

The overall difference is estimated correctly by the *N*_3_ and *L*_2_*N* methods automatically, and accounting for *θ*_0_ *≠*= 0 yields results which are practically identical to the simulations in which *θ*_0_ = 0 in terms of power and FDR. In contrast, when we use limma the FDR is no longer controlled at the desired level (0.05, in these simulations.) For example, with *α* = 9, *θ* ∈ {2.5, 3}, *D* = 250, and there is a small overall mean shift, *θ*_0_ = 0.5, limma’s actual FDR is 0.11. When the overall mean shift is greater, *θ*_0_ = 1, limma’s actual FDR is 0.39.

To ensure that the FDR is controlled at the desired level when using the limma modeling approach in DVX, one has to center the gene expression data around the group means (or medians) for each gene. That is, if *y*_*ijg*_ if the expression level of gene *g* for subject *j* in group *i*, transforming it to 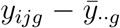 resolves the problem of high false positive rate. The DVX software provides an easy way to transform gene expression data, and in particular it allows the user to perform median-centering.

### 3.4 Case Study

Understanding the mechanisms that preserve normal neuronal functionality is very important for treating Alzheimer’s disease (AD) patients. REST/NRSF (repressor element 1-silencing transcription/neuron-restrictive silencer factor) is known to regulate neuronal genes during embryonic development, and Lu et al. [15] showed that it is “induced in the aging human brain and regulates a network of genes that mediate cell death, stress resistance and AD pathology.” Lu et al. [15] observed that REST is lost from the nucleus of cells among AD and mild cognitive impairment (MCI) patients, which leads to dysregulation of this gene network.

In the experiment, gene expression levels for 54,675 genes were obtained from 41 people, in four groups: extremely aged (95-106yr) (n=4), normal aged (70-94yr) (n=16), middle aged (40-69yr) (n=9), and young (*<*40yr) (n=12). There are 21 females and 20 males in this sample. The data has been deposited in the National Center for Biotechnology Information (NCBI) Gene Expression Omnibus (GEO) repository with accession number GSE53890.

The deposited data has been normalized, but we observed some skewness which can be explained by a large number of low-abundance and low-variance genes. We eliminate these genes, as they may be indistinguishable from ‘background noise’. We filter out genes which have an overall median log-expression *≤*5.5, across all subjects. We also equalized the medians across all samples, in order to ameliorate any subject-specific effects. The resulting dataset contained 17,833 genes. Since the change in neuronal condition is known to deteriorate gradually over time for adults, we tested three contrasts: young vs. middle aged, middle aged vs. normal aged, and normal aged vs. extremely aged. We performed the differential analysis using *L*_2_*N, N*_3_ and limma, and in all cases we controlled for gender.

Figure 2 shows the fitted distributions for each of the three contrasts, namely, Young as the baseline group vs. Middle-aged as treatment (left), Middle-aged vs. Normal-aged (center), and Normal-aged vs. Extremely aged (right). Sub-figures A-C were obtained by fitting the *L*_2_*N* model, and D-F were obtained by using limma. The red curve represents the distribution of the ‘null’ (non differentially expressed) genes, and the green curves show the distribution of the non-null genes, per the selected model. Note that limma, by default, uses P(non-null) = 0.01. The dashed blue line is the fitted mixture distribution. These plots also show the estimated goodness of fit, in terms of the root mean squared error (rMSE). For all three contrasts, the *L*_2_*N* model yields a better fit than limma. For example, rMSE=0.02 vs. 0.35 for the Young vs. Middle-aged comparison. Note that the scales on the x-axis are different for limma and *L*_2_*N* (and *N*_3_, which is not shown here.) For limma, *dE* is the estimated contrast between the two groups, accounting for predictors. If no predictors are included in the model, *dE* is just the difference between the mean expression level in the treatment group and that in the control group. When using *N*_3_ or *L*_2_*N* to fit the data, the x-axis is labeled *dE*_*v*_ which is the estimated standardized contrast between the two groups, accounting for predictors. The standardized contrasts are obtained by dividing *dE* by the estimated gene-specific standard deviations.

**Figure 2:**
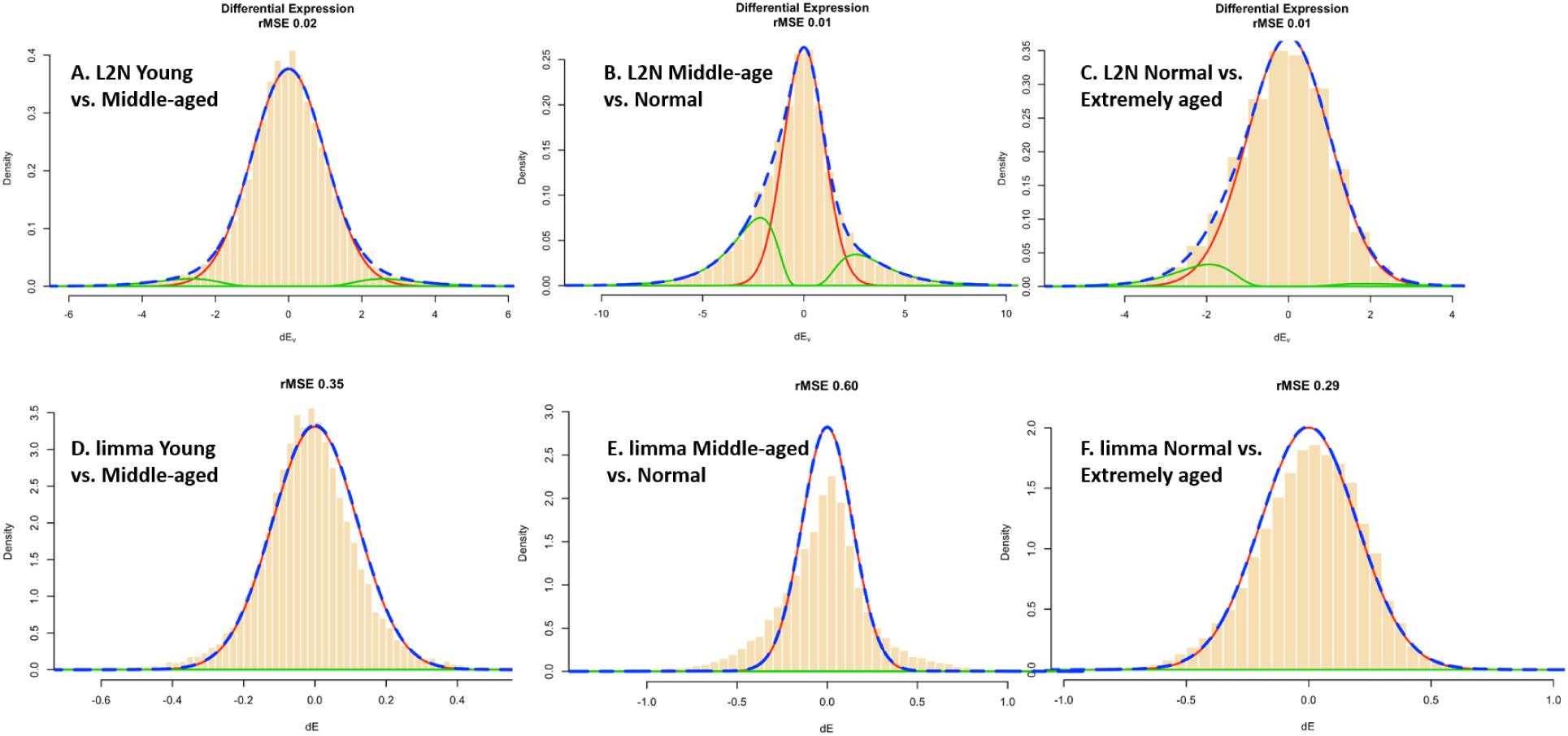
Fitted distributions for the differential expression statistics. The sub-figures in the top row (A-C) were obtained by fitting the *L*_2_*N* model. Sub-figures D-F were obtained by using limma. Panels A and D depict the fitted distributions for the comparion between middle aged and young (the baseline); B and E correspond to the comparison between Middle-aged (baseline) and Normal-aged, and C and F correspond to the contrast Normalaged (baseline) vs. Extremely-aged.

It is clear from the plots that in the comparisons young vs. middle aged and normal aged vs. extremely aged, the vast majority of genes are not differentially expressed, whereas in the comparison between middle aged vs. normal aged many genes are estimated to be differentially expressed. Furthermore, plots B and C show that according to the *L*_2_*N* model, there are more genes which are overexpressed in the younger cohort (the baseline group in each comparison), since the mixture component in the (*-8*, 0] range has a higher weight than the component in the [0, *8*) range.

Both *L*_2_*N* and *N*_3_ also test for differential variation. With this dataset and with a q-value [19] threshold of 0.01, no genes are differentially dispersed in any of the three comparisons. The number of differentially expressed genes obtained from each method for each of the three contrasts (using q *≤*0.01) is summarized in Table 4. The columns labeled “B*>*T” (”T*>*B”) contain the total number of genes found to be overexpressed in the baseline (treatment) group. The numbers in parentheses are the root mean squared errors for the fitted model. In all three comparisons, *L*_2_*N* gives the best fit, in terms of the rMSE. In the comparison between Young and Middle Aged, no genes are found to be DE when using limma, while *N*_3_ and *L*_2_*N* find 157 DE genes with q-value *≤* 0.01. Similarly, in the comparison between “Normal Aged” and “Extremely Aged” limma detects no DE genes, and *N*_3_ and *L*_2_*N* find 64 DE genes. All three methods detect hundreds of DE genes in the comparison between “Middle Aged” and “Normal Aged” (limma: 522, *N*_3_: 2,682, *L*_2_*N*: 2,725), suggesting that a significant change, as pertains to cognition, takes place after 70yrs.

**Table 4:**
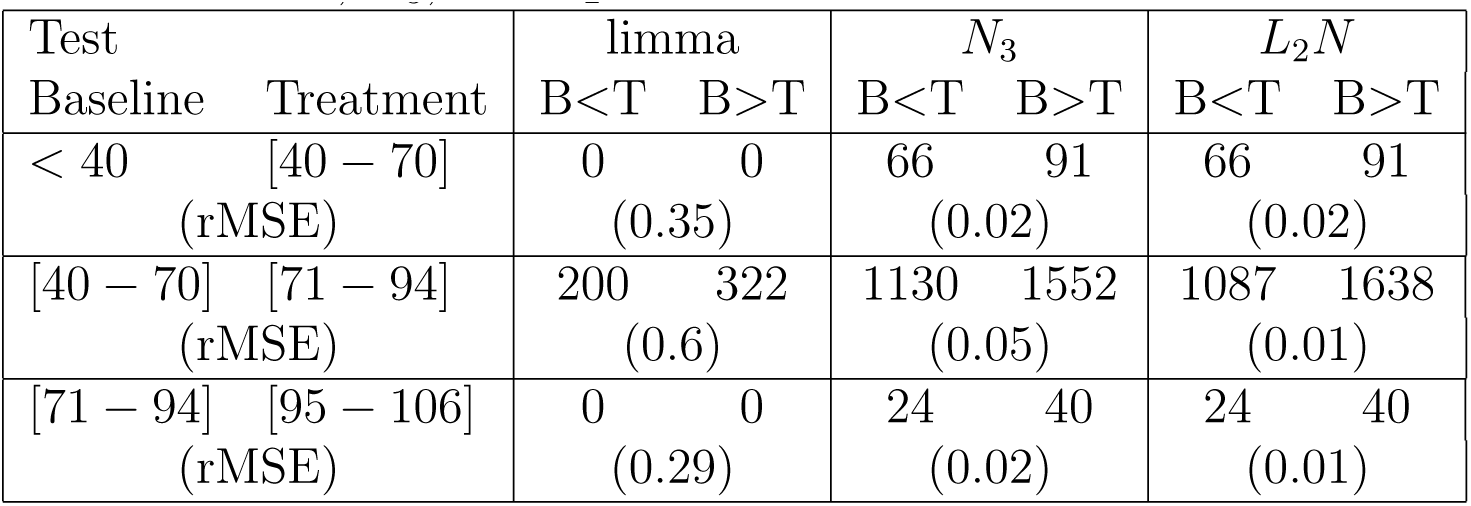
Case study - the REST dataset (GSE53890) - total number of differentially expressed genes obtained from limma, *N*_3_, and *L*_2_*N*.

A detailed version of this case study which demonstrates several features of the DVX package (e.g., use of diagnostic plots and data transformation tools, generation of report and result files in Microsoft Word- and Excel-compatible formats, respectively) is available in the supplementary material.

## 4 Discussion

We introduced the *L*_2_*N* three-component mixture model and corresponding empirical Bayes implementation as a flexible approach for assessing differential variation and expression. The proposed model is particularly well-suited for situations in which there is no a priori knowledge of the mixture distribution of the data: it can capture asymmetries in the numbers of over-expressed [dispersed] and under-expressed [dispersed] genes when they exist and can provide a better fit to null and non-null components that are not well-separated, while still performing well under symmetry and little overlap. In our power analysis, we compared *L*_2_*N* with two other hierarchical mixture models, namely, limma [18], and *N*_3_ which consists of three normal distributions [2]. A key feature common to all three models is that the differential expression *d*_*g*_ of non-DE genes are assumed to come from a normal distribution. Each model assumes that the DE genes arise from a common distribution, but the choice of nonnull distribution differs across the three models. This hierarchical modeling approach results in ‘borrowed information’ across genes, which leads to greater power, when compared to naive (one gene at a time) approaches.

Both *L*_2_*N* and *N*_3_ have two advantages over limma. First, the DE genes are allowed to have a non-symmetrical prior distribution, which makes the three-component mixture models less restrictive, and more realistic in many cases, as demostrated by the case study. Second, the same mixture model can be used to account for differential variation when evaluating differential (mean) expression. We see in our simulations that this leads to improved power when differential variation truly exists.

*L*_2_*N* was shown to be a bit more powerful than *N*_3_ when differential dispersion exists, and more so when the variability of the random errors increases. Furthermore, in our simulations and analysis of real datasets, *L*_2_*N* seems to provide the best fit to the differential expression statistics in terms of rMSE. Also, of the three mixture models, *L*_2_*N* is the only one which uses non-local priors for the DE genes. Conceptually, this is advantageous because it implies that there is a negligible probability of declaring a gene as DE when the actual differential expression statistic is close to 0.

To make the method easily accessible to biologists, we created a user-friendly R Shiny interface called DVX [4], which also includes the implementations of *N*_3_ and limma. With all three models, it is possible to control for the effects of other factors and covariates, and to set up linear contrasts, beyond the simple two-group comparison. DVX offers a set of diagnostic plots and transformation tools, and options for exporting Word-compatible reports and Excel-compatible result tables. DVX may also be used with count data (e.g., RNA-seq read counts) with proper data transformation, using tools such as ‘voom’ (see, e.g., [14] and [13].)

## Acknowledgements

We wish to thank Ved Deshpande and M. Henry Linder for their help with earlier versions of the code and documentation.

